# EM-Compressor: Electron Microscopy Image Compression in Connectomics with Variational Autoencoders

**DOI:** 10.1101/2024.07.07.601368

**Authors:** Yicong Li, Core Francisco Park, Daniel Xenes, Caitlyn Bishop, Daniel R. Berger, Aravi D.T. Samuel, Brock Wester, Jeff W. Lichtman, Hanspeter Pfister, Wanhua Li, Yaron Meirovitch

## Abstract

The ongoing pursuit to map detailed brain structures at high resolution using electron microscopy (EM) has led to advancements in imaging that enable the generation of connectomic volumes that have reached the petabyte scale and are soon expected to reach the exascale for whole mouse brain collections. To tackle the high costs of managing these large-scale datasets, we have developed a data compression approach employing Variational Autoencoders (VAEs) to significantly reduce data storage requirements. Due to their ability to capture the complex patterns of EM images, our VAE models notably decrease data size while carefully preserving important image features pertinent to connectomics-based image analysis. Through a comprehensive study using human EM volumes (H01 dataset), we demonstrate how our approach can reduce data to as little as 1/128th of the original size without significantly compromising the ability to subsequently segment the data, outperforming standard data size reduction methods. This performance suggests that this method can greatly alleviate requirements for data management for connectomics applications, and enable more efficient data access and sharing. Additionally, we developed a cloud-based application named EM-Compressor on top of this work to enable on-thefly interactive visualization: https://em-compressor-demonstration.s3.amazonaws.com/EM-Compressor+App.mp4.

## 1 Introduction

The field of connectomics aims to create detailed, synapse-level maps of neural circuits, promising to deepen our understanding of the brain’s structural complexities. These maps are indispensable for elucidating the neural circuit functions underlying behavior, but obtaining them is highly resource-intensive. Connectomics datasets require the generation of high-resolution image volumes and segmentations that reach petabyte scale for a cubic millimeter [25] and exabyte scale for a whole mouse brain [1]. Consequently, the field faces imminent data storage bottlenecks within processing and analysis pipelines, necessitating innovative data storage and processing solutions.

Recent advancements in imaging and image processing have started to confront these data scale challenges, from rapid and intelligent ML-based acquisition of large serial section scanning EM (ssSEM) datasets [19,24] to cost-effective image analysis algorithms [18,14,15]. Ideally, these methods would enable a seamless integration of image analysis processes such as segmentation within the image acquisition step, reducing the need for temporary data storage or extensive data transfer. However, current practices separate these modules by location (across different facilities for imaging and computation) and time (months to years from acquisition to segmentation). As a result, laboratories require petabyte-scale data storage and sharing solutions, which are not commonly affordable for most research teams.

We thus identify a clear need and opportunity to simplify data management for connectomics: improved compression of EM data to reduce data storage and transfer costs. While current image compression methods in the computer vision field may be suitable for EM image data, an idealized solution must not only reduce data volume size but also preserve the quality and fidelity of image features crucial for accurate automated neural segmentation. Traditional image compression algorithms like JPEG [29], JPEG2000 [5], and AVIF^3^ are efficient to compute, but suffer from blocking artifacts and color bleeding when generating compressed images, especially when the target compression rate is high. With the rapid development of Deep Learning, image processing and computer vision researchers have proposed various learning-based algorithms [27,28,26,2,4,8,20,21] to address challenges for image compression. However, most of these methods are evaluated on natural image datasets, and their performance, especially the ability to maintain segmentation quality on reconstructed images after compression, remains unknown when applied to EM image data, which differ in appearance and content. The closest prior work focused on related challenges addressed here is [22], which presents a solution tailored for EM data by first denoising and then applying standard codecs such as AVIF, achieving a compression rate of 17 times. Inspired by this work, we intend to push the boundary of the compression rate significantly using end-to-end learning-based methods, towards addressing data bottlenecks in the current connectomics pipeline, a major challenge for the field. Additionally, since connectomic data analysis relies on neuroscientists’ ability to navigate large datasets in real-time, we investigate the feasibility of rapid, on-the-fly encoding and decoding in cloud services for current data exploration tools [16,3,11,30] to enable real-time visualization and proofreading.

Herein we provide a comparative analysis assessing the performance of a modified version of the original VAE [13] to compress human brain [25] EM images against standard codecs such as JPEG2000 and AVIF. Our main contributions are as follows:

- To our best knowledge, we are the first to apply learning-based methods to EM-based and connectomics-focused image compression by designing and training multiple VAE models with different middle feature sizes covering a wide range of compression rates. This work provides insights into the architecture design choice of VAE as well as its strengths and limitations in compressing connectomic image data, improving the compression rate from 17 times [22] to 128 times, outperforming standard image compression codecs without significantly compromising the ability to segment the data.
- We conducted evaluations of the compression results, assessing the ability to retain domain-specific image content needed for downstream tasks in connectomics research by testing on neuron membrane and instance segmentation.
- We developed a cloud-based application named EM-Compressor to enable on-the-fly compression of EM images. We integrate this application with a commonly used visualization tool called Neuroglancer [16], enabling neuroscientists to interactively inspect the reconstructed images after compression.

## 2 Method

### 2.1 Workflow of EM-Compressor

Fig. 1 shows the overall workflow of the proposed EM-Compressor. The input to the compressor is a set of stitched EM images of brain tissue. We use the encoder part from the trained Variational Autoencoder (VAE) model to compress the images to the feature space. The elements in the feature vector are originally represented in 32-bit format and can be transformed into 16-bit, 8-bit, and 4-bit formats for higher compression rates before being sent to storage. During the decoding process, we apply the decoder of the trained VAE model to the stored features to reconstruct “original” EM images for potential downstream analysis tasks such as neuron instance segmentation. The whole process except for image acquisition is wrapped into a cloud-based application.

**Fig. 1.**
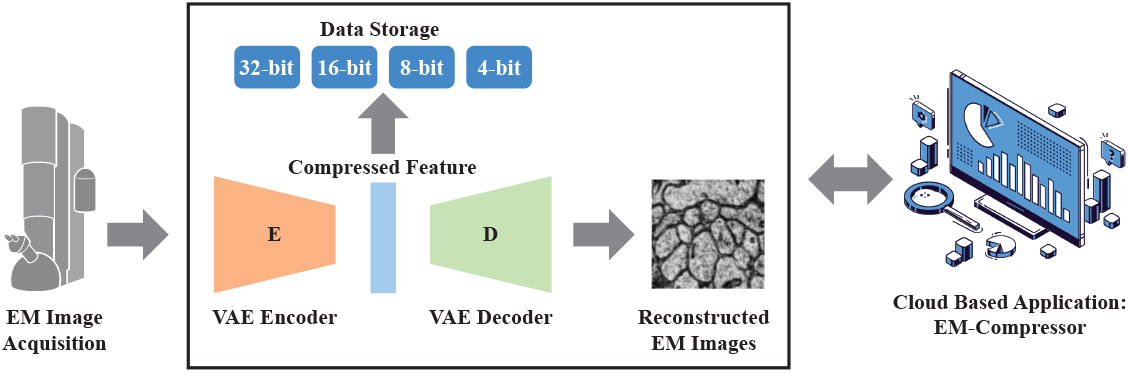
Workflow of EM-Compressor. Connectomic images are first acquired by EM. Next, the encoder of VAE is applied to compress EM images to a feature vector, whose elements are represented in 32-bit, 16-bit, 8-bit, and 4-bit formats for storage. To reconstruct the image from the feature, the decoder of VAE is applied to perform upsampling. This data compression process is wrapped into a cloud-based application.

### 2.2 Variational Autoencoder

The 2D convolutional VAE consists of an encoder and a decoder; the former downsamples the input image, and the latter upsamples the stored features. Each layer in the encoder contains two consecutive ResNet [9] blocks, and spatial downsampling by a factor of 2 in each dimension. Similarly, each layer in the decoder spatially upsamples by a factor of 2 in each dimension. Specifically, given an image *x* with size of *H × W*, by passing it through the encoder *E*, we get a feature *f* = *E*(*x*) with a size of 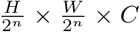, where n is the number of downsample layers and C is the number of channels in the VAE’s middle feature space, depending on the target compression rate. Suppose *x* is represented in *i*-bit format and we want to further reduce the data size by representing *f* in *j*-bit format, the compression rate *r* we can get is:

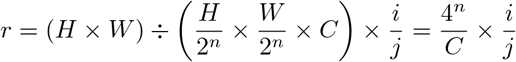

During the decoding process, we pass *f* through the decoder *D* to get the recon-structed image 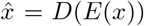 with the same size as input image *x*. The overall loss *L*_*t*_ for training our VAE consists of three parts as follows:

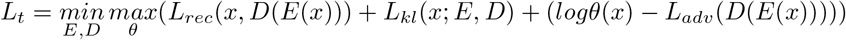

where the first term *L*_*rec*_ is a pixel-wise loss such as *L*_1_ or *L*_2_, the second Kullback-Leibler-term *L*_*kl*_ is to prevent the model from creating a high variance latent space by penalizing the latent to be zero-centered with small variance, similar to the original VAE [13]. Training on a pixel-wise loss often leads to blurry images, therefore we add the third term which is an adversarial loss following [6,7,12] to enforce that the reconstructed image matches the visual quality of the uncompressed input image. *θ* is a discriminator that tries to distinguish the reconstructed image from the uncompressed image.

## 3 Experiments

### 3.1 Dataset

For all experiments in this paper, we use the publicly-available H01 dataset [25], containing a 1.4-petabyte image volume of human brain tissue acquired by ssSEM, with its corresponding neuron membrane and instance segmentation (license^4^). To train the VAE models with good generalization, we cropped 10 subvolumes from well-separated regions with distinct tissue appearances and split them for training (7 subvolumes), validation (1 subvolume), and testing (2 subvolumes). Each subvolume has 2048 2D image sections with a size of 512*×*512.

### 3.2 Evaluation

We assess the compression performance of all methods using segmentation criteria, i.e., Dice and Intersection over Union (IoU) for neuron membrane segmentation and Variation of Information (VI)^5^ [17] for neuron instance segmentation. The motivation is that while metrics like MSE can evaluate the performance of a compression algorithm, this metric is agnostic of downstream connectomics processing: generating accurate dense segmentation of individual neurons. However, it is critical for a compression model to retain the ability to delineate individual cellular objects within each image, even at the cost of losing information that may not be relevant for dense segmentation. To this end, we trained a model for membrane and instance segmentation on uncompressed images and compared the performance of uncompressed and compressed images across different compression methods and rates.

Specifically, to assess the ability to label the individual neuronal objects (instance segmentation), we use VI to provide a topological measure of the merge and split errors occurring in the image data due to compression. The VI between two image labels *S*_1_ and *S*_2_ can be calculated by comparing the partition entropy of each image with the cross labeling *S*_1_ *× S*_2_. VI=VImerge+VI where

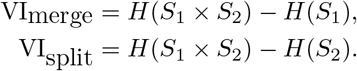

### 3.3 Setup

#### Image compression

We compress EM images in JPEG2000 format with Glymur^6^ covering the following compression rates: 2x, 4x, 8x, 16x, 32x, 64x, and 128x. For AVIF format, we use Pillow^7^‘s AVIF plugin and cover the following compression rates: 2x, 4x, 8x, 16x, 32x, and 64x. 128x is not reachable in AVIF format. As for the image compression via VAE, we trained 12 models, 2 for each starting compression rate (one with bigger spatial size and one with more channels in terms of the middle feature size). For example, a model denoted “vae 16x 1×64×64” indicates a starting rate of 16x compression with a middle feature size of 1*×*64*×*64, and by further changing the original 32-bit representation to 16/8/4-bit we can achieve 32x, 64x, and 128x. Therefore, 12 models with 6 starting compression rates (2x, 4x, 8x, 16x, 32x, 64x) enable us to compress from 2x to 512x. All VAE models are trained for 50 epochs on 4 NVIDIA A100 GPUs with a batch size of 2 for each. The best checkpoint is selected with the validation set.

#### Segmentation model

To evaluate each compression method’s performance on membrane and instance segmentation task, we trained a U-Net[23] with GELU activation[10] with training data sampled at random locations and angles also from the H01 dataset [25]. The network was trained for 50000 gradient steps with the cross entropy loss and the Adam optimizer with *lr* = 0.01.

#### Cloud-based deployment for EM-Compressor

EM-Compressor visualization application is hosted on an AWS g4dn.16xlarge instance, which features an Nvidia T4 GPU, 64 vCPU, and 256GB of internal memory. The application is powered by Streamlit, a Python-based web framework, and content is served under an application load balancer.

## 4 Results

### 4.1 Quantitative comparison

Fig. 2 demonstrates a quantitative comparison of VAE-based methods against the JPEG2000 and AVIF codecs on the neuron membrane and instance segmentation tasks at different compression rates. According to the five metrics (IoU, Dice, VI-Total, VI-Split, VI-Merge), we find out that VAE-based methods are comparable with JPEG2000 and AVIF at relatively low compression rates (up to 8x). As the target compression rate increases, VAE-based methods significantly outperform the two standard codecs. Additionally, since our 12 VAE models have results at overlapping compression rates, we observe that in most cases, training a model that has a large feature size followed by minimizing the bit representation to reach a certain compression rate has better performance than directly training a model that has a small feature size at that same compression rate. From an architecture design perspective, comparing the two VAE models with the same starting rate, in most cases, we notice that the model with a middle feature that has a larger spatial size is better than the model with a middle feature that has a larger number of channels, e.g, “vae_32x_2×32×32” is better than “vae_32x_8×16×16”. We also observe the unsatisfactory results achieved by both vae 64x models, due to the limitation of our method in the sense that more downsampling layers make it harder to maintain the reconstruction quality.

**Fig. 2.**
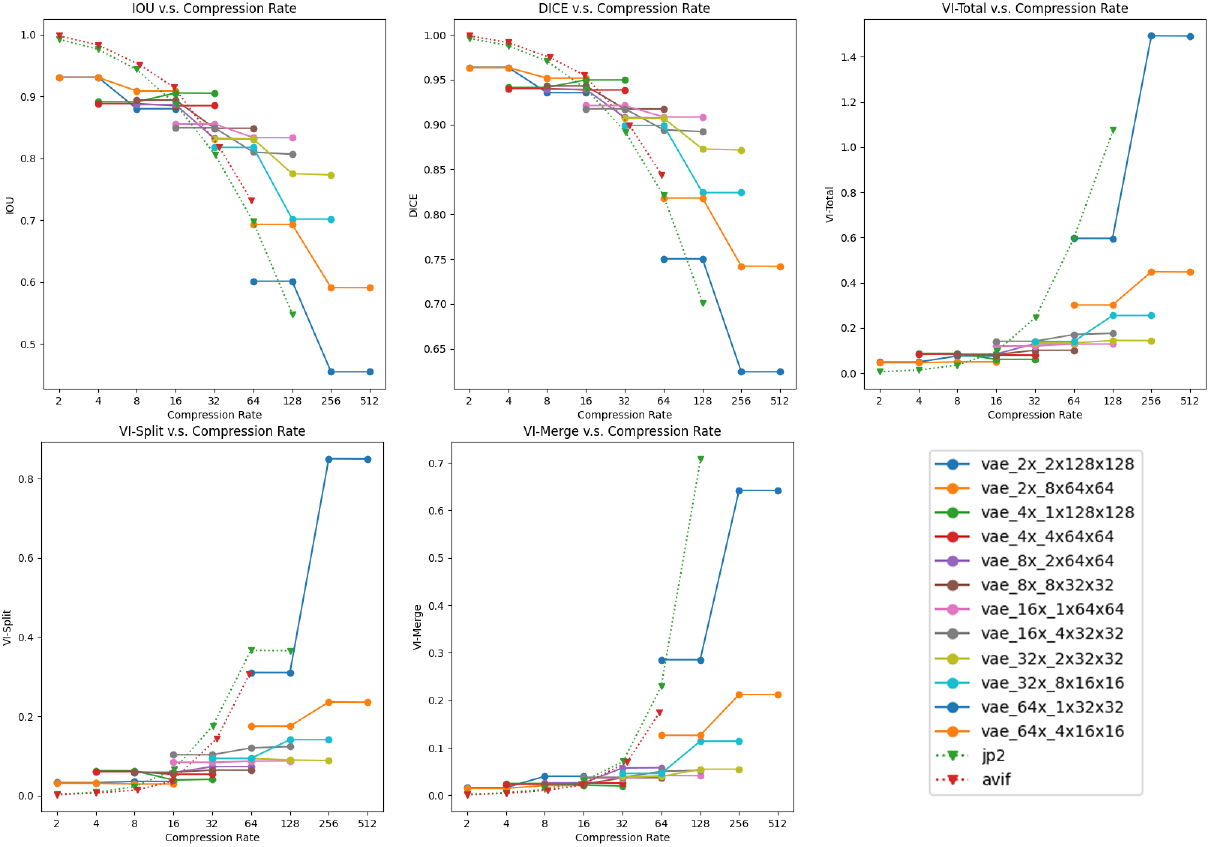
Quantitative comparison of different compression methods on the neuron membrane and instance segmentation task at different compression rates with segmentation metrics including IoU, Dice, and VI. For relatively low compression rates (up to 8x), AVIF and JPEG2000 perform comparably to or slightly better than VAE-based methods. As the compression rate gets larger, VAE-based methods perform significantly better than standard codecs. The results are obtained on the test set described in Section 3.1.

### 4.2 Qualitative comparison

We further conducted a qualitative comparison of different compression methods on EM images. Fig. 3 combines both the neuron membrane and instance segmentation results. We evaluated our method alongside AVIF and JPEG2000 at compression rates ranging from 16x to 64x. We observe that our method exhibits superior performance compared to AVIF and JPEG2000 in several key aspects. First, as the compression ratio increases, a noticeable degradation in image quality is observed in AVIF and JPEG2000, characterized by increased noise, heightened blurring, and pronounced blocking artifacts. In contrast, our approach effectively preserves the original image quality to a satisfactory degree. Second, our approach effectively preserves the intricate patterns and structures inherent in EM images that are important for connectomics research. At each compression rate, our method demonstrates fewer merge and split errors, thereby yielding more accurate instance segmentation outcomes. These findings underscore the efficacy of our approach in maintaining the visual quality of EMs, which is essential for accurate connectomics image analysis.

**Fig. 3.**
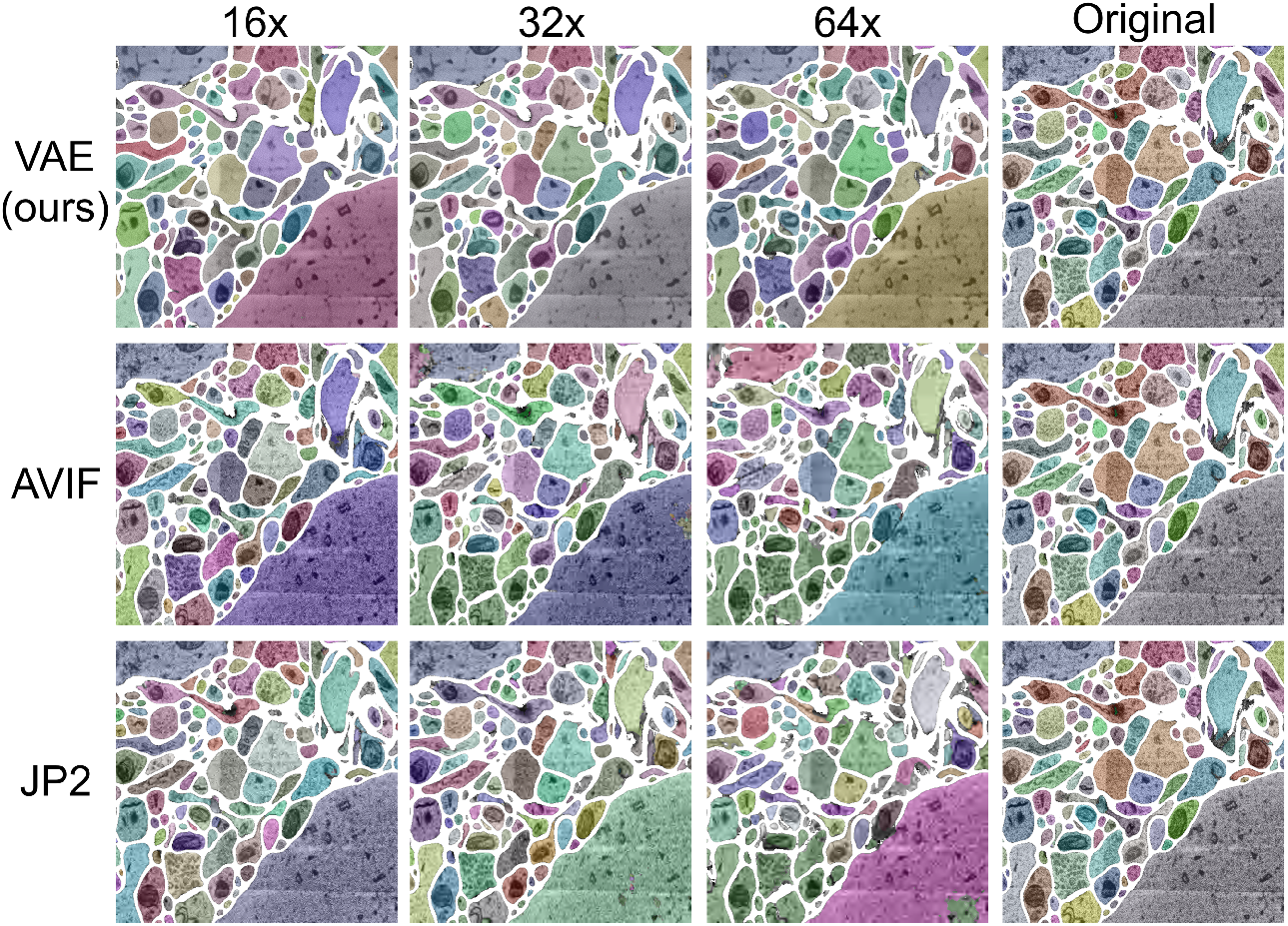
Qualitative comparison of different compression methods on the neuron membrane and instance segmentation task at 16x, 32x, and 64x compression rates. In comparison to the original uncompressed image, results from our VAE-based method maintain good segmentation quality, whereas AVIF and JPEG2000 increasingly suffer from merge and split errors as the compression rate increases. The results are obtained on the test set described in Section 3.1.

### 4.3 Cloud-based application: EM-Compressor

To facilitate broader access and interactive exploration, we developed a publicly available cloud-based application called EM-Compressor based on the trained VAE models. EM-Compressor provides two core features: single image compression comparison and batch precomputed compression. The former is best suited for real-time 2D comparison across VAE-based and traditional compression methods and ratios with a self-provided image. The latter feature enables compression and visual comparison with uncompressed 3D image data through Neuroglancer [16]. The application is hosted on a cloud instance, ensuring highly available, on-demand testing and execution of EM-Compressor without requiring users to possess high-end computational resources. A video demonstration is available at https://em-compressor-demonstration.s3.amazonaws.com/EM-Compressor+App.mp4.

## 5 Discussion and Conclusion

Connectomic data generation and processing is experiencing rapid advancements, accelerated by improvements in machine learning and the facilitation of cloud-based resources. Owing to these advances, data storage and transfer are emerging as a growing bottleneck necessitating new approaches for data compression. By leveraging machine learning, specifically through learned dimensionality reduction with VAEs, our EM-Compressor data compression approach has achieved a reduction in EM image data size of over two orders of magnitude without significantly affecting image quality or features necessary for connectomics-related data processing. We also developed a cloud-based interactive visualization tool based on our method. In addition, we discovered in VAE feature space, larger spatial resolution are generally more favorable than larger number of channels in terms of high-fidelity neural segmentation. Future improvements could include leveraging 3D data or optimizing VAE models for segmentation tasks to allow higher compression rates without quality loss. However, we note that volumetric VAEs may not suit all connectomics applications due to the need for near-perfect 3D alignment.

## Footnotes

3 https://aomediacodec.github.io/av1-avif/

4 https://creativecommons.org/licenses/by-nc-nd/4.0/

5 https://pypi.org/project/python-voi

6 https://glymur.readthedocs.io/en/latest/

7 https://pillow.readthedocs.io/en/stable/

